# Genomes to Fields 2026: Genomic resources for investigating genotype-phenotype relationships in *Zea mays*

**DOI:** 10.64898/2026.11.15.766802

**Authors:** Nicholas A. Johnson, Ann Murithi, Corrinne E. Grover, Shawn Kaeppler, Dayane Cristina Lima, Hirva Bhayani, Dinuri Lokuwalpola, Heidi Chen, Nicholas Allsing, Todd P. Michael, Edward S. Buckler, Matthew B. Hufford, Maria C. Romay

**Affiliations:** Institute for Genomic Diversity, Cornell University, 130 Biotechnology Building, NY 14853 USA; Department of Ecology, Evolution, and Organismal Biology, Iowa State University, Ames, IA 50011 USA; Wisconsin Crop Innovation Center, University of Wisconsin – Madison, 8520 University Green, Middleton, WI 53562 USA; Department of Plant and Agroecosystem Sciences, University of Wisconsin – Madison, 1575 Linden Drive, Madison, WI 53706 USA; The Plant Molecular and Cellular Laboratory, Salk Institute for Biological Studies, La Jolla, CA, 92037, USA; USDA-ARS Plant, Soil and Nutrition Research Unit, RW Holley Center, 538 Tower Rd, Ithaca, 14853 NY USA

## Abstract

Maize (*Zea mays* L.) is a major global crop characterized by high genetic and phenotypic diversity across accessions. To better understand this diversity, the Genomes-to-Fields initiative has produced paired genotypic and phenotypic datasets annually since 2014. Here, we present 15 genome assemblies representing 13 maize lines, including founders and testers from the 2022–2023, 2024–2025, and 2026–2027 Genomes-to-Fields evaluation cycles and additional breeding germplasm, together with gene and repeat annotations. Assemblies ranged from 2.19 to 2.38 Gb, with BUSCO completeness of 97.9–99.0%. Excluding the more fragmented LH287 assembly (contig N50 = 5.3 Mb; 90.8% chromosome placement), contig N50 values ranged from 31.1 to 195.8 Mb, with 95.9–99.4% of sequence placed on chromosomes. Gene annotation identified approximately 40,000 gene models per genotype, of which 93.6–93.9% were classified as protein-coding, with 95.9–97.3% complete BUSCOs. Comparison with B73 identified 27,606–30,517 one-to-one syntenic orthologs per genotype, representing 69.4–76.8% of B73 genes. Transposable elements occupied 83.6–88.6% of the genomes, with LTR retrotransposons representing the predominant class. Comparative genome alignments revealed three large inversions, including a 12.65-Mb inversion on chromosome 3 present in six of the 13 lines. Together, these resources expand the genomic representation of maize breeding germplasm and provide a foundation for investigating genetic variation underlying agronomic traits.

## Introduction

Maize (*Zea mays* L.) is one of the most widely cultivated crops globally, providing a major source of food, animal feed, and industrial raw materials. Beyond its agricultural significance, maize is a model system for plant genomics and evolution, with a transposable element-rich genome and extensive genetic and structural variation among accessions (Hufford et al., 2021; Schnable et al., 2009). This diversity, shaped by adaptation and breeding, presents opportunities for understanding the genetic basis of complex agronomic traits and their responses to heterogeneous growing environments.

Recent advances in long-read sequencing have enabled increasingly complete maize genome assemblies, revealing extensive structural variation that cannot be fully represented by a single reference genome. Comparative analyses of the maize nested association mapping (NAM) founder genomes have highlighted substantial differences in gene content, chromosomal structure, and repetitive sequences (Hufford et al., 2021). Such variation can have important functional consequences. Chromosomal inversions can suppress recombination and maintain linked haplotypes, and have been associated with differences in flowering time, leaf morphology, and yield-related traits in maize and its wild relatives (Crow et al., 2020; Fustier et al., 2019; Romero Navarro et al., 2017). Similarly, heterochromatic knobs, composed of tandem repeats, can influence local recombination and, in certain genetic backgrounds, chromosome transmission, while their repetitive structure presents challenges for genome assembly (Ghaffari et al., 2013; Kanizay et al., 2013; Liu et al., 2020). Despite substantial advances in characterizing maize genomic diversity, much remains to be discovered about the extent of pangenomic variation and its contribution to agronomically important traits across environments.

The *Genomes to Fields* (G2F; https://www.genomes2fields.org/home/) initiative is a public–private partnership established in 2013 to advance understanding of how maize genetic variation influences phenotype across diverse environments. Through multidisciplinary collaboration, G2F integrates publicly available genomic, phenotypic, and environmental data with computational modeling to characterize genotype-by-environment (G×E) interactions, improve predictions of agronomically important traits, and inform crop improvement (Chen et al., 2026; Lima et al., 2023).

Here, we present genomic resources comprising long-read sequencing data, 15 genome assemblies representing 13 maize lines, and associated gene and repeat annotations. These resources encompass founder, tester, and additional breeding germplasm from the 2022–2023, 2024–2025, and 2026–2027 G2F evaluation cycles. Comparative analyses identified three large chromosomal inversions, assessed gene model correspondence with B73, and characterized transposable element content across these genomes. Together, these resources support future investigations of maize genomic diversity and genotype–phenotype relationships within G2F.

## Methods and materials

### Germplasm selection

Four stiff-stalk inbred lines (‘3AZA1’, ‘FBLL’, ‘PHB47’, and ‘PHBW8’) were crossed with four stiff-stalk-derived Germplasm Enhancement of Maize (GEM) lines: ‘GEMS-0061’, ‘GEMS-0113’, ‘GEMS-0219’, and ‘GEMS-0227’ (Rogers et al., 2022). Doubled-haploid (DH) lines were derived from the resulting F1 progeny and subsequently testcrossed with the ex-Plant Variety Protection (ex-PVP) inbred lines ‘PHP02’ and ‘LH287’ for the 2024–2025 evaluation cycle (Figure 1; https://www.genomes2fields.org/about/project-overview/). Three additional lines represent other evaluation cycles: ‘LH244’, a stiff-stalk tester used in 2022–2023, and ‘PH09B’ and ‘PH2MW’, ex-PVP inbreds included in the 2026–2027 evaluation cycle. The pedigree, origin, and distinguishing characteristics of each line are described below, based on records from the USDA National Plant Germplasm System (GRIN-Global), Plant Variety Protection (PVP) documentation, and published germplasm reports (National Plant Germplasm System, 2025; White et al., 2020).

**Figure 1.**
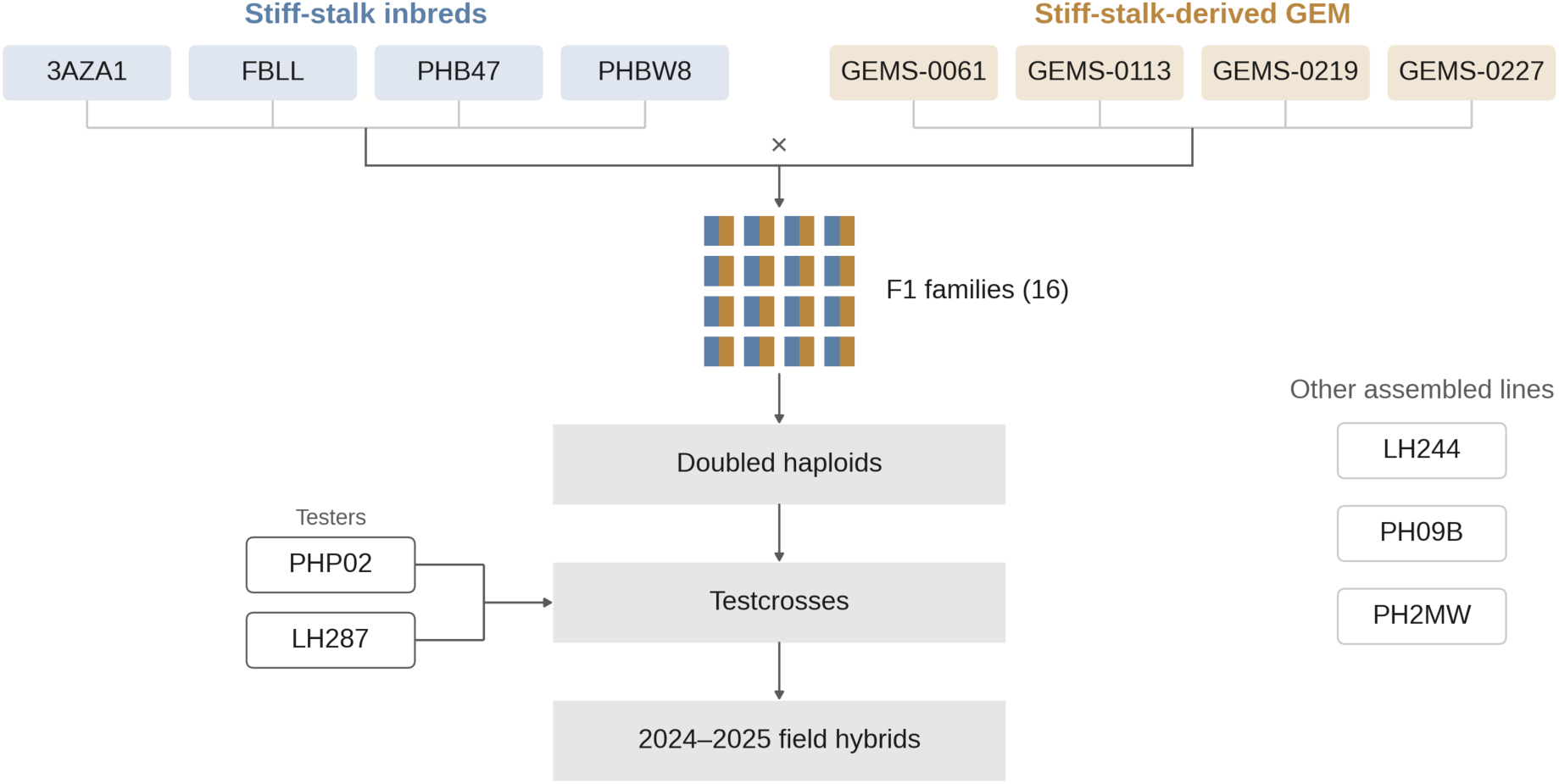
Breeding scheme and germplasm selection for the Genomes-to-Fields maize population. Four stiff-stalk inbred lines (3AZA1, FBLL, PHB47, and PHBW8) were crossed with four stiff-stalk-derived Germplasm Enhancement of Maize (GEM) lines (GEMS-0061, GEMS-0113, GEMS-0219, and GEMS-0227), generating 16 F1 families. Doubled-haploid (DH) lines derived from these crosses were testcrossed with PHP02 and LH287 to produce hybrids for the 2024–2025 field evaluation cycle. Three additional assembled lines represent other evaluation cycles: LH244 (2022–2023) and PH09B and PH2MW (2026–2027).

### 3AZA1

Former DEKALB Genetics Corporation PVP (PVP 9600027) maintained by the USDA National Plant Germplasm System (GRIN-Global) under PI 592379. This is a highly inbred stiff stalk pedigree line developed from proprietary germplasm incorporating several public inbreds. 3AZA1 was developed by crossing AQA3 X QFAR, with AQA3 derived from A632, CM105, B14, and LH23, and QFAR derived from B14, B37, and LH23 (USDA GRIN-Global, PI 592379). Its characteristics include yellow semident kernels, red cobs, and approximately 16 cm ears.

### FBLL

Former DEKALB Plant Genetics PVP-associated inbred (PVP 9100034) maintained by GRIN-Global under PI 546481. This is a highly inbred, yellow dent line with a breeding background related to B73-type germplasm. FBLL was developed by crossing the proprietary DEKALB lines 5B2C-A X PB80 in 1982, followed by multiple generations of self-pollination and selection, USDA Plant Inventory No. 200 (Pfund, 2002). Its characteristics include yellow dent kernels, red cobs, approximately 14.4 cm ears, and an average of 17 kernel rows per ear.

### GEMS-0061

Germplasm Enhancement of Maize (GEM) breeding line maintained by GRIN-Global under PI 685801. This is a partially inbred, exotic-derived line developed through the USDA-ARS GEM project using South American maize germplasm. GEMS-0061 was developed from the Argentine accession ARZM 16 026 (PI 516027), originating from Mendoza, crossed with the anonymous stiff stalk GEM donor S17, with the selected pedigree AR16026:S17-10-1-B-B (USDA GRIN-Global, PI 685801). Its characteristics include predominantly yellow to orange-red kernels exhibiting semident and semiflint morphology, white cobs, and approximately 15.5 cm ears.

### GEMS-0113

Germplasm Enhancement of Maize (GEM) breeding line maintained by GRIN-Global under PI 651540. This is a partially inbred, exotic-derived line released through the USDA-ARS GEM project in 2005. GEMS-0113 was developed from the Mexican accession CHIS775 (PI 515164) and the S1911b breeding donor, with the selected pedigree CHIS775:S1911b-120-001-B-B-SIB (USDA GRIN-Global, PI 651540). Its characteristics include yellow semident kernels, red cobs, regularly arranged kernel rows, and approximately 14–15 cm ears.

### GEMS-0219

Germplasm Enhancement of Maize (GEM) breeding line maintained by GRIN-Global under PI 685914. This is a partially inbred, exotic-derived line developed through the USDA-ARS GEM project and released in 2011. GEMS-0219 was developed from the Cuban maize accession CUBA164 (PI 489361) and GEM breeding line GEMS-0002 (PI 632413), with the selected pedigree (CUBA164:S2012-459-001-B/GEMS-0002)-B-B-031-B-B-SIB (USDAGRIN-Global, PI 685914). Its characteristics include yellow semident kernels, white cobs, approximately 13–14.5 cm ears, and plant heights of approximately 242 cm in a USDA 2016 evaluation.

### GEMS-0227

Germplasm Enhancement of Maize (GEM) breeding line maintained by GRIN-Global under PI 685920. This is a partially inbred, exotic-derived line developed through the USDA-ARS GEM project and released in 2012. GEMS-0227 was developed from the tropical maize accession NEI9004 (Ames 29045), associated with Suwan germplasm, with the selected pedigree NEI9004:S2809-B-022 (USDA GRIN-Global, PI 685920). Its characteristics include yellow semident kernels, red cobs, approximately 14 kernel rows, and 15.5 cm ears.

### LH244

Former Holden’s Foundation Seeds inbred (PVP 200000122) maintained by GRIN-Global under PI 612589. This is a highly inbred stiff stalk line developed from Holden’s breeding germplasm and used as a tester in the Genomes to Fields 2022/2023 evaluation cycle. LH244 was developed by crossing LH197 X (LH197 X LH199), representing a backcross toward LH197 (USDA GRIN-Global, PI 612589). Its characteristics include yellow dent kernels, red cobs, approximately 14–16 kernel rows, and 15.5–17 cm ears. LH244 is also notable for its high transformation efficiency and extensive genomic similarity to B73 (Tan et al., 2026).

### LH287

Former Monsanto inbred line (PVP 200000121, maintained by GRIN-Global as PI 612588) developed via a cross between LH212Ht X LH185 (Foley, 2001). LH287 is a yellow dent, non-stiff stalk line that is widely used as a tester in hybrids, including in the Genomes to Fields initiative (https://www.genomes2fields.org/about/project-overview/). Characteristics of LH287 include pollen shed at 70 days post emergence, yellow-orange dent kernels, white cobs and 16 cm ears.

### PH09B

Former Pioneer Hi-Bred PVP (PVP 9700218) maintained by the USDA National Plant Germplasm System (GRIN-Global) under PI 597331. This is a high-yielding, highly inbred stiff stalk line frequently used in commercial hybrids with PH2MW. This accession was developed by crossing PHP38 X PHHB9, PVP Certificate (USDA Plant Variety Protection Office, 2000), and its characteristics include pollen shed at approximately 74 days post emergence, yellow dent kernels, red cobs, and 17-19 cm ears.

### PH2MW

Former Pioneer Hi-Bred PVP (PVP 9900382) maintained by GRIN-Global as PI 612160. This is a non-stiff-stalk line historically used as a pollen parent in hybrid production. PH2MW was developed by crossing PHGG6 X PHAW6, PVP Certificate (USDA Plant Variety Protection Office, 2001), and its characteristics include pollen shed at 76 days post emergence, yellow dent kernels, red cobs, and 18 cm ears.

### PHB47

Former Pioneer Hi-Bred PVP (PVP 8300141, registered as B47) maintained by GRIN-Global under PI 601009. This is a highly inbred stiff stalk line derived from public maize breeding germplasm. PHB47 was developed by crossing B37 X SD105, with B37 originating from Iowa Stiff Stalk Synthetic and SD105 representing an early public South Dakota inbred (USDA GRIN-Global, PI 601009). Its characteristics include yellow kernels exhibiting semident to semiflint morphology, red cobs, and approximately 14–16 cm ears in multiple USDA evaluations.

### PHBW8

Former Pioneer Hi-Bred PVP (PVP 9200079) maintained by GRIN-Global under PI 559936. This is a highly inbred stiff stalk line developed from Pioneer breeding germplasm. PHBW8 was developed by crossing PHJ40 X PHW52 (USDA GRIN-Global, PI 559936). Its characteristics include yellow semident kernels, red cobs, approximately 14 kernel rows, and ears measuring 18 cm in a USDA 2014 evaluation.

### PHP02

Former Pioneer Hi-Bred PVP (PVP 8800212) maintained by GRIN-Global under PI 601570. This is a highly inbred line used as a tester in commercial maize breeding and in the Genomes to Fields initiative. PHP02 was developed by crossing PHG44 X PHG29 (USDA GRIN-Global, PI 601570). Its characteristics include yellow dent kernels, red cobs, and approximately 14.5 cm ears.

### Genome sequencing and assembly

PacBio HiFi reads were generated on the Revio system and processed on-instrument using ccs v8.0.1, trim v1.1.0, and lima v2.10.0. HiFi BAM files were converted to FASTQ using bam2fastq (pbtk v3.5.0; https://github.com/PacificBiosciences/pbtk/tree/v3.5.0). Oxford Nanopore Technologies (ONT) reads (R10.4.1 chemistry, PromethION 2 Solo) were basecalled using MinKNOW with the HAC v4.2.0 model (SUP v4.2.0 for PHP02). Reads passing MinKNOW quality filtering and ≥50 kb in length were selected for hybrid assembly using SeqKit v2.13.0 (Shen et al., 2016).

Read-set identities were assessed using k-mer fingerprinting (*k* = 21 and 31) with sourmash v4.9.3 (Irber et al., 2024). Potential sample discordance was further investigated by examining haplotype structure across individual SMRT cells, allele concordance between sequencing technologies, and the alignment of reads to their corresponding assemblies.

Genome assemblies were generated using hifiasm v0.25.0-r726 (Cheng et al., 2021, 2024), incorporating HiFi and ONT ultra-long reads where appropriate. Assembly configurations were selected based on read concordance, haplotype structure, assembly completeness, sequence duplication, and structural consistency. For six lines (FBLL, PHBW8, PHB47, 3AZA1, PHP02, and GEMS-0227), unpurged primary contigs (-l0) were retained. For GEMS-0061 and GEMS-0113, partially phased assemblies were generated using haplotig purging (-l3) with adjusted primary coverage parameters (--pri-range 24 and 23, respectively), and both haplotype partitions were retained. Evidence of mixed biological material in GEMS-0219 prompted assembly from HiFi reads originating from a single SMRT cell (bc2041), using-l3 --pri-range 16 without ONT reads and retaining only the primary assembly.

### Assembly quality control

Assembly contiguity statistics were calculated using SeqKit. Gene-space completeness was evaluated using BUSCO v6.1.0 (Tegenfeldt et al., 2025) in genome mode with miniprot and the poales_odb12.2 lineage dataset (6,188 orthologs). B73 RefGen_v5 (Hufford et al., 2021) was evaluated using the same procedure for comparison. Consensus quality values (QV) and k-mer completeness were estimated using Merqury v1.3 (Rhie et al., 2020) with *k* = 21 databases generated from the corresponding HiFi reads.

Assembly integrity was evaluated using read alignments, assembly graphs, and synteny with B73 RefGen_v5. HiFi reads were aligned to their respective assemblies using minimap2 v2.30; map-hifi (Li, 2018), with alignments processed using SAMtools v1.20 (Danecek et al., 2021) to assess sequence coverage and structural support. Gene-anchor synteny relative to B73 was evaluated using AnchorWave v1.3.1 (Song et al., 2022). Structural differences from B73 were not considered assembly errors without additional supporting evidence. Six unsupported interchromosomal joins were resolved by sequence-preserving contig splits, including two contradicted by read evidence and four associated with unsupported joins within (AGT)n repeat arrays.

All 11 contig assemblies were screened for foreign-organism contamination with NCBI FCS-GX v0.5.5 (Astashyn et al., 2024) against the GX database (build 2023-01-24) using taxonomy ID 4577 (Zea mays), and for adaptor and vector sequence with FCS-adaptor v0.5.5 in eukaryotic mode (--euk).

### Reference-guided scaffolding

Contigs were then assigned to chromosomes, ordered, and oriented relative to B73 RefGen_v5 using ALLMAPS JCVI v1.6.6 (Tang et al., 2015). A maize pan-genome anchor-marker set comprising 100-bp sequences was mapped to B73 to establish reference positions. Markers were subsequently aligned to the assembled contigs using HISAT2 v2.2.1 (Kim et al., 2019), retaining uniquely mapped alignments (MAPQ 60; edit distance ≤5). Contigs of at least 100 kb containing at least 10 unique markers at five distinct reference positions were eligible for placement.

Scaffolds were constructed using ALLMAPS, with a maximum of 100 evenly distributed markers per contig. Adjacent contigs were joined using 100-N gaps of unknown physical length, without trimming or splitting contig sequences. Scaffold joins were evaluated using HiFi read alignments, assembly-graph connectivity, and ONT reads where available from the same biological sample.

ONT reads from GEMS-0061 and GEMS-0113 were excluded from join validation because of uncertain biological concordance with the HiFi samples. Joins were classified according to direct sequence support or reference-based placement, and conflicting evidence was individually reviewed.

Contigs lacking sufficiently supported chromosome assignments were retained as unplaced sequences. Scaffold sequences were validated against their AGP representations to confirm that each contig was incorporated completely and without duplication. Because chromosome assignments and contig order were inferred using B73, the resulting scaffolds represent reference-guided chromosome organization rather than independently resolved chromosome-scale assemblies.

Telomeric repeat arrays were identified with tidk v0.3.2; motif TTTAGGG;-w 10000 --min-length 200 --max-offset 1000 (Brown et al., 2025), and chromosome ends were classified from these arrays. The Mo17 CAU-2.0 telomere-to-telomere (T2T) assembly and B73 RefGen_v5 served as controls.

The assembly pipeline was implemented as a containerized Snakemake workflow v9.26.1 (Mölder et al., 2021), with accompanying documentation and AI-agent instructions to facilitate reproducible execution and maintenance.

### Genome annotation

Genomes were annotated using Helixer 0.3.3, which leverages a cross-species deep learning model for accurate *ab initio* gene structure prediction (Holst et al., 2026). The model identifies genomic regions corresponding to the untranslated regions (UTRs), coding sequences (CDSs), and introns of genes. For each species, one of four predefined lineage-specific models (vertebrate, invertebrate, land plant, or fungi) is selected based on its taxonomic classification. Here, for maize, the land plant lineage model was used.

### Gene model annotation validation

The completeness of gene annotations was evaluated with Benchmarking Universal Single-Copy Orthologs (BUSCO v6.1.0) in protein mode, using the Poales lineage dataset (poales_odb12), which contains 6,282 conserved single-copy orthologs. For each of the genomes, the predicted proteins were searched against the lineage dataset, and the numbers of complete (single-copy and duplicated), fragmented, and missing BUSCOs were recorded. To assess whether the predicted gene models are likely to encode proteins, every Helixer-predicted gene model was scored with Protein Sequence Assessment Using a Reference ORF Network (PSAURON v1.1.3) (Sommer et al., 2025). PSAURON is a machine-learning model trained on annotated protein-coding sequences to estimate the probability that a CDS encodes a protein, both in its annotated reading frame (in-frame score) and in the five alternative frames (out-of-frame scores). Extracted CDS sequences were run on PSAURON using default settings, and gene models with an in-frame score ≥0.5 and out-of-frame scores ≤0.5 were classified as true protein-coding sequences.

### Orthologous genes between each genome and B73

Syntenic genes between each of the genomes and B73 (RefGen_v5) were identified with MCScan (JCVI utility libraries v1.6.8) (Tang et al., 2024). B73 gene model and the gene models of each genotype were aligned with LAST (Kiełbasa et al., 2011), and the parameters “--cscore=.99 --quota=1:1” were used to find 1-to-1 syntenic gene relationships. A similar approach was used to identify syntenic genes between A188 and B73 (Yao et al., 2026). To validate this approach, Liftoff was also used to lift over all genomes to the B73 (RefGen_v5) coordinate system to identify genes that matched B73 genes. The resulting lifted GFF3 files were processed to extract CDS features, and the overlapping CDS regions between the B73 reference genes and transferred gene models from each of the genomes were identified using BEDTools intersect (Quinlan & Hall, 2010). The cross-referenced, corresponding gene identifiers were extracted from the resulting intersection files using the Pandas package in Python (V3.14.10). For both analyses, a candidate orthologous gene matrix was constructed using B73 genes as the reference with 1-to-many gene relationships for liftoff and 1-to-1 relationship for MCScan.

### Transposable element annotation

Transposable elements (TEs) were annotated across the genomes using EDTA v2.3.0 (Ou et al., 2019) with maize-specific settings (--species Maize). The pipeline identified different TE classes using LTRharvest (Ellinghaus et al., 2008), LTR_FINDER (Xu & Wang, 2007), TIR-Learner (Su et al., 2019), HelitronScanner (Xiong et al., 2014) AnnoSINE2 v2.0.9, and the EDTA LINE module. The TE libraries were combined and curated using EDTA, accompanied by de novo predictions from RepeatModeler v2.0.7 (Flynn et al., 2020), and used by RepeatMasker v4.2.3 (http://www.repeatmasker.org) to annotate intact and fragmented TE copies. The genomic coverage of each TE class was then calculated across genomes.

### AI usage

Claude Code (Opus 5.5; v2.1.318; Anthropic) was used to generate and audit code. ChatGPT (GPT-6; OpenAI) was used for reviewing and editing writing and to audit Claude Code output.

## Results and discussion

### Sequencing and genome assembly

Genome assemblies for thirteen maize lines were generated using accession-specific sequencing and assembly strategies, including PacBio HiFi and Oxford Nanopore (ONT) long-read sequencing. Among twelve lines with available sequencing records, HiFi inputs ranged from 57 to 150 Gb (25–65× coverage), while ONT reads ≥50 kb contributed 0.03–1.95 Gb (less than 1× coverage; Supplementary Table 1). Differences in residual heterozygosity and sequencing data quality required accession-specific assembly methods, resulting in 11 primary assemblies and two partially phased assembly pairs.

The 11 primary assemblies ranged from 2.19 to 2.38 Gb, with contig N50 values of 94.9–195.8 Mb for 10 assemblies, while LH287 was substantially more fragmented (N50 = 5.3 Mb). The partially phased haplotype partitions (2.24–2.30 Gb) had contig N50 values of 31.1–85.8 Mb (Table 1). Assembly completeness was high, with 97.9–99.0% of conserved BUSCO genes recovered and estimated consensus QV values of 67.5–76.0. Primary assemblies recovered 95.8–98.7% of read-derived k-mers, while the partially phased haplotype pairs collectively recovered approximately 98%. Across all 15 assemblies, three contigs spanned more than one chromosome (two unresolved terminal joins in GEMS-0227 and one misjoin in PH2MW), telomeric repeat arrays ≥1 kb were detected at 249 of 300 chromosome ends (83.0%), and 12 chromosomes were assembled as single gap-free contigs with telomeric arrays at both ends. To facilitate downstream comparative analyses, assemblies were organized into chromosome-assigned scaffolds using accession-specific approaches, placing 95.9–99.4% of assembled sequence (90.8% for LH287) into scaffolds assigned to the 10 maize chromosomes. Because some intercontig joins were supported by reference order alone, these scaffolds should be interpreted as chromosome assignments rather than uniformly contiguous chromosome-level assemblies.

**Table 1:**
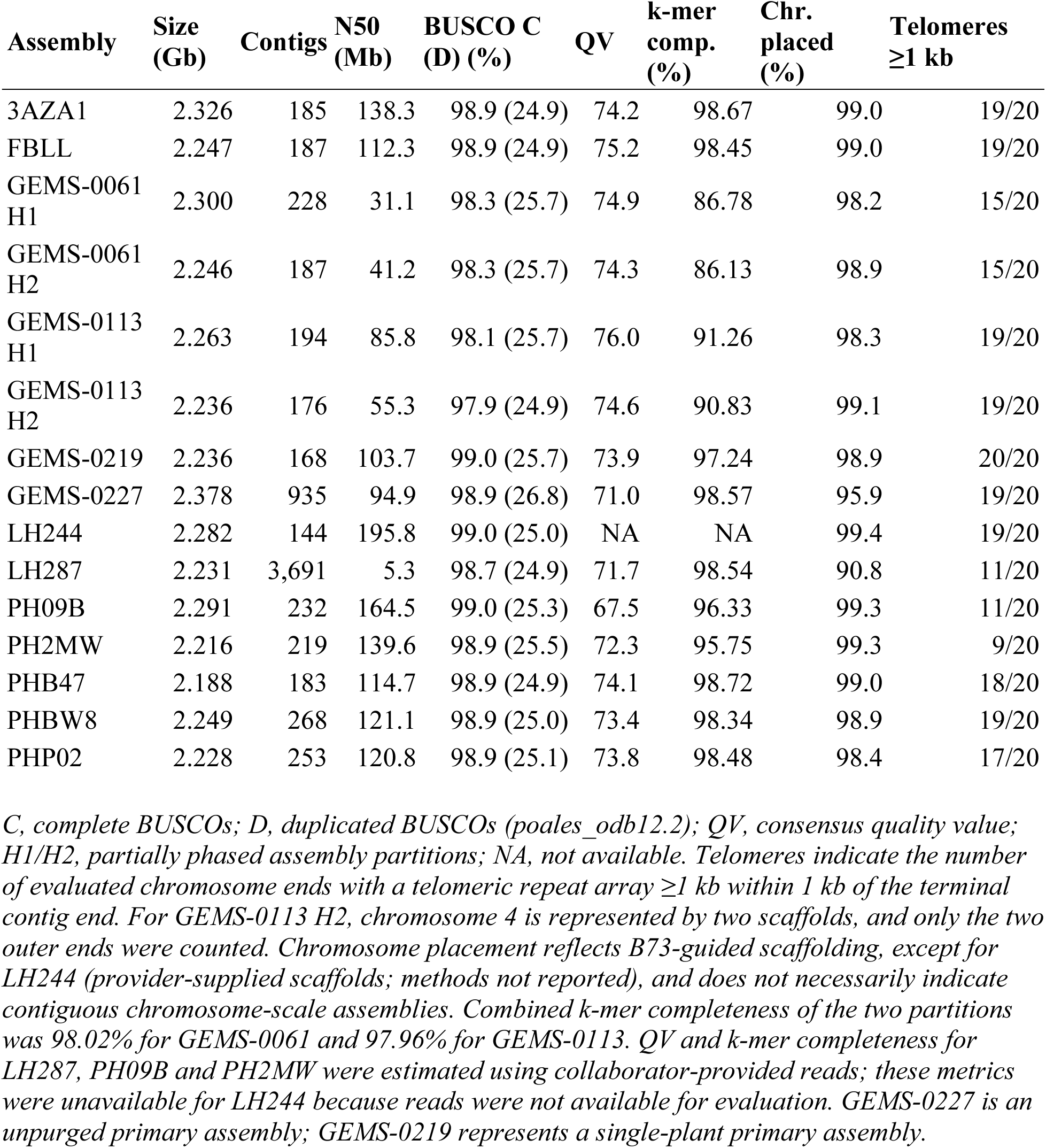
Assembly statistics and quality metrics for thirteen maize lines.

### Genomic structural variation

Comparative alignments with B73 RefGen_v5 revealed three well-supported large inversions across the 13 maize lines (Figure 2). Inversion arrangements were identified from assembled sequences spanning both breakpoints or, for the fragmented LH287 assembly, from long reads supporting both inversion junctions.

**Figure 2:**
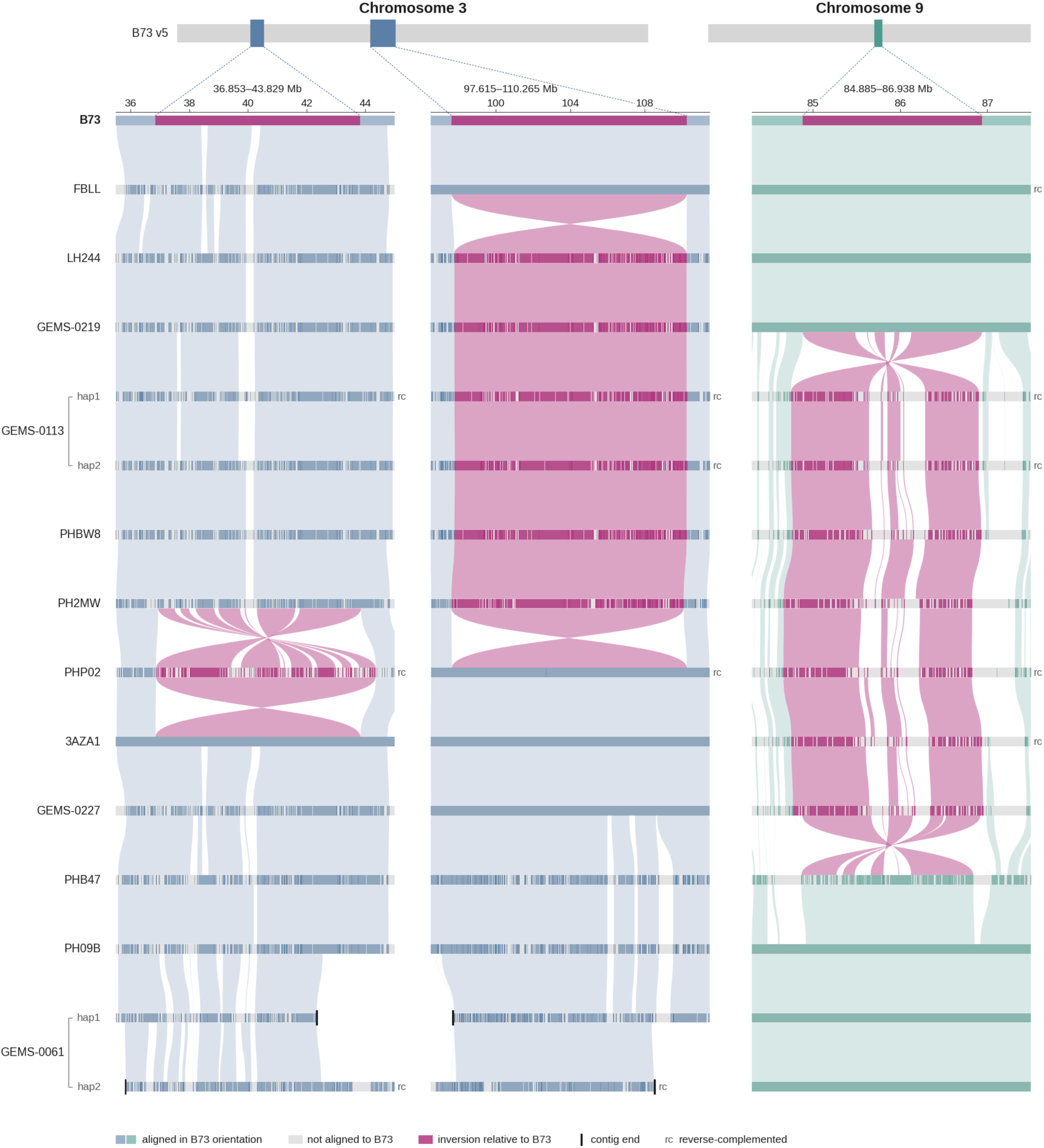
Chromosomal inversions in G2F 2026 maize assemblies. B73 v5 ideograms indicate two regions on chromosome 3 (36.853–43.829 and 97.615–110.265 Mb) and one on chromosome 9 (84.885–86.938 Mb), enlarged below (dashed lines). Assembly contigs are shown at their original lengths, aligned to B73. Blue (chromosome 3) and green (chromosome 9) indicate sequences in B73 orientation, magenta indicates sequences aligned in reverse orientation relative to B73, and grey indicates regions without ordered B73 alignments. Ribbons connect corresponding sequences, with crossings indicating orientation differences. Black bars mark contig ends; ‘rc’ indicates reverse-complemented contigs; and hap1/hap2 denote the two partially phased partitions of GEMS-0061 and GEMS-0113. LH287 is not shown because its assembly is fragmented at these loci.

The largest inversion spanned 12.65 Mb on chromosome 3 (97.6–110.3 Mb) and was present in six of 13 lines (GEMS-0113, GEMS-0219, LH244, LH287, PH2MW and PHBW8). Breakpoints were resolved to base-pair precision and were identical among assembled carriers. In five lines, 17–39 HiFi or ONT reads per junction supported the inverted arrangement, with no qualifying reads supporting the B73 orientation; LH244 was supported by assembly alignments alone. Both junction sequences were also present in the genome assemblies of NAM founders P39 and Oh43 (Hufford et al., 2021), and corresponded to inversion SV3 previously reported in LH244 (Tan et al., 2026).

A second inversion spanning 2.05 Mb on chromosome 9 (84.9–86.9 Mb) was identified in seven of 13 lines (3AZA1, GEMS-0113, GEMS-0227, LH287, PH2MW, PHBW8 and PHP02), with 23–54 orientation-switching long reads supporting each junction in all seven lines. A 6.98-Mb inversion on chromosome 3 (36.9–43.8 Mb) was found only in PHP02 among the 12 lines that could be assessed, with more limited read support at both junctions; the same junction sequences were present in the NAM founder CML228 assembly (Hufford et al., 2021). The two partially phased partitions of GEMS-0061 and GEMS-0113 did not disagree at any of the three loci. Overall, nine of the 13 lines carried at least one of these inversion polymorphisms, demonstrating the established structural variation known to exist across maize lines.

### Gene annotation and completeness

Similar numbers of genes were annotated in each genome. Helixer predicted between 39,868 (GEMS-0113 hap2) and 41,074 (PH2MW) gene models across the genotypes (Table 2). Each gene model had a single transcript, so mRNA and gene counts were identical. The average CDS lengths were ∼1,139 bp across the genotypes, with the longest average found in line PHBW8 at 1,145 bp and the shortest average found in line GEMS-0061 hap2 at 1,134 bp. Overall, gene feature statistics, including mean CDS length, exon and intron lengths, protein length, and exon number, were highly consistent across the genotypes (Figure 3; Supplementary Table 2). BUSCO analysis reported 95.9–97.3% complete BUSCOs across the genotypes. Complete BUSCOs comprised 70.1–72.3% single-copy and 24.8–27.0% duplicated genes (Table 2; Supplementary Table 3). PSAURON, on the other hand, classified 93.6–93.9% of the gene models in each genotype as protein-coding (Table 2), with a mean in-frame score of 0.906–0.909 (Supplementary Table 4). Low-scoring gene models (in-frame score < 0.5) were more common among genes with shorter CDS lengths, and their count decreased as CDS length increased. Single-exon gene models also had a higher proportion of low scores than multi-exon models across all genotypes (Supplementary Table 4).

**Figure 3:**
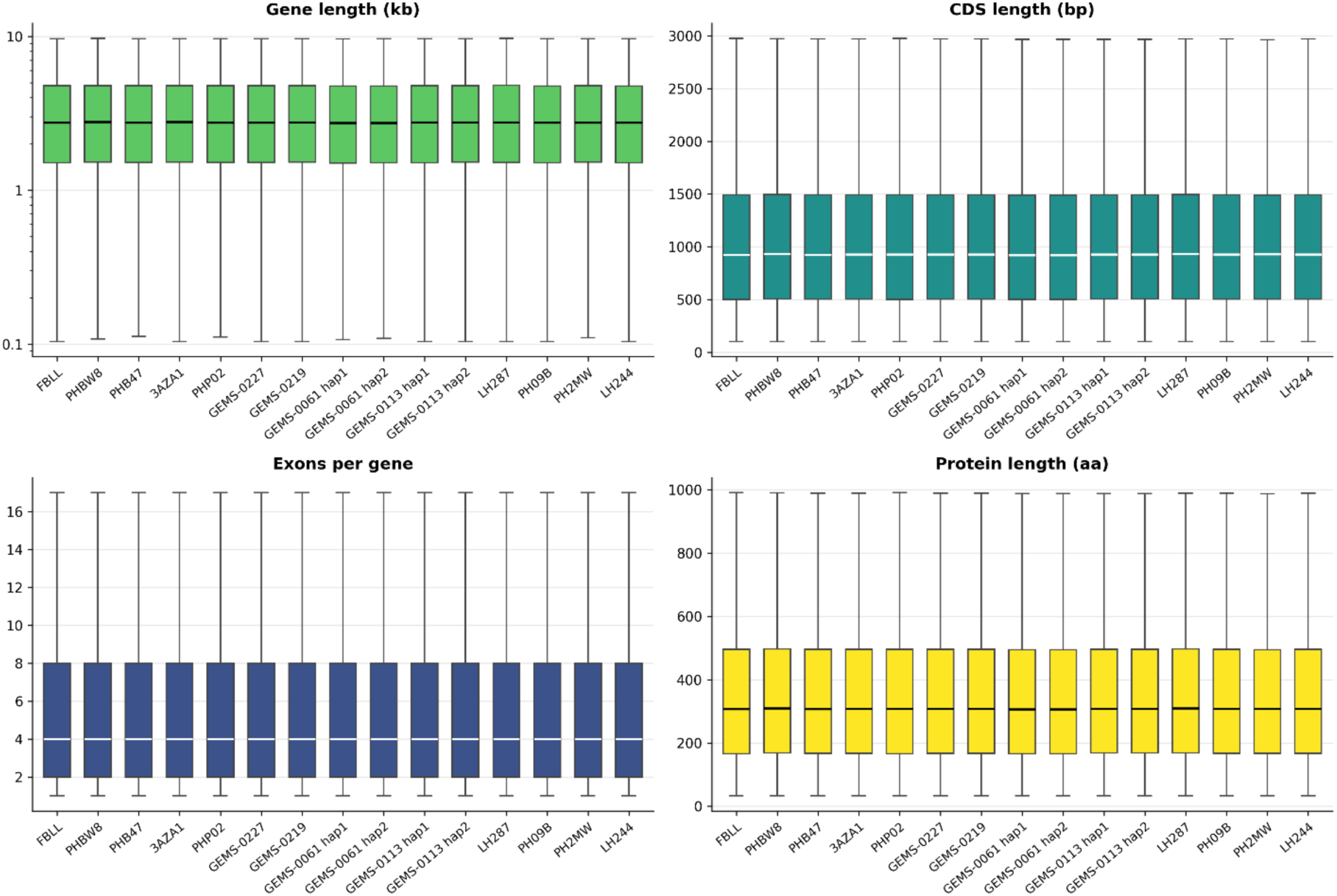
Comparison of gene model characteristics across 13 maize genotypes. Distributions of gene length (kb), coding sequence (CDS) length (bp), exon number per gene, and predicted protein length (amino acids) across annotated gene models. Boxplots show the median, interquartile range, and whiskers. Gene model characteristics were broadly consistent across genotypes.

**Table 2:**
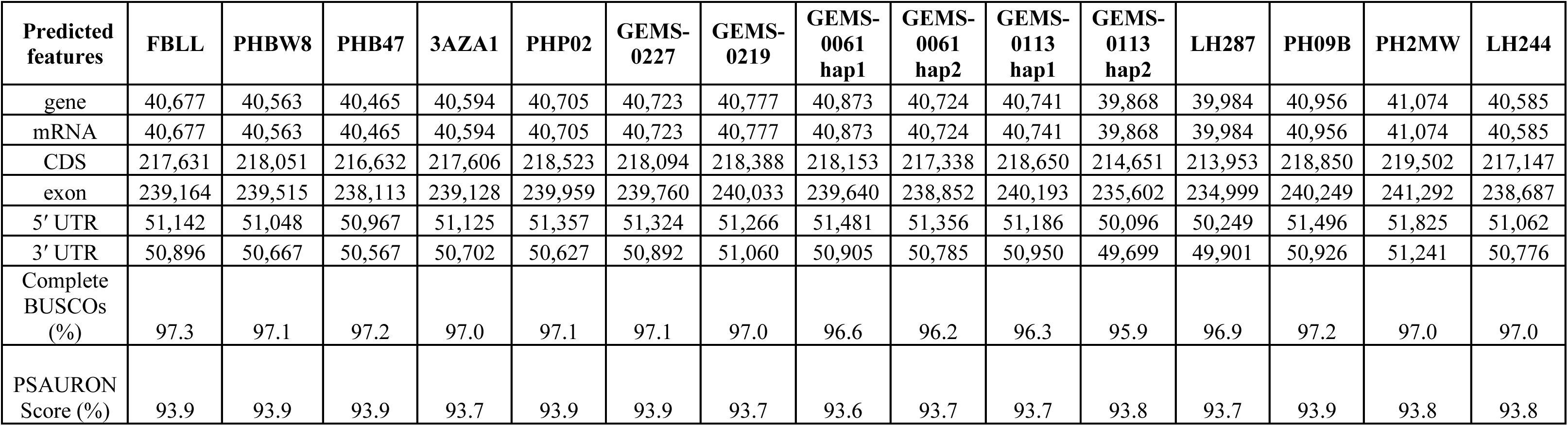
Summary statistics for the gene features predicted by Helixer on each genome assembly.

| **Predicted features** | **FBLL** | **PHBW8** | **PHB47** | **3AZA1** | **PHP02** | **GEMS-0227** | **GEMS-0219** | **GEMS-0061 hap1** | **GEMS-0061 hap2** | **GEMS-0113 hap1** | **GEMS-0113 hap2** | **LH287** | **PH09B** | **PH2MW** | **LH244** |
|---|---|---|---|---|---|---|---|---|---|---|---|---|---|---|---|
| gene | 40,677 | 40,563 | 40,465 | 40,594 | 40,705 | 40,723 | 40,777 | 40,873 | 40,724 | 40,741 | 39,868 | 39,984 | 40,956 | 41,074 | 40,585 |
| mRNA | 40,677 | 40,563 | 40,465 | 40,594 | 40,705 | 40,723 | 40,777 | 40,873 | 40,724 | 40,741 | 39,868 | 39,984 | 40,956 | 41,074 | 40,585 |
| CDS | 217,631 | 218,051 | 216,632 | 217,606 | 218,523 | 218,094 | 218,388 | 218,153 | 217,338 | 218,650 | 214,651 | 213,953 | 218,850 | 219,502 | 217,147 |
| exon | 239,164 | 239,515 | 238,113 | 239,128 | 239,959 | 239,760 | 240,033 | 239,640 | 238,852 | 240,193 | 235,602 | 234,999 | 240,249 | 241,292 | 238,687 |
| 5′ UTR | 51,142 | 51,048 | 50,967 | 51,125 | 51,357 | 51,324 | 51,266 | 51,481 | 51,356 | 51,186 | 50,096 | 50,249 | 51,496 | 51,825 | 51,062 |
| 3′ UTR | 50,896 | 50,667 | 50,567 | 50,702 | 50,627 | 50,892 | 51,060 | 50,905 | 50,785 | 50,950 | 49,699 | 49,901 | 50,926 | 51,241 | 50,776 |
| Complete BUSCOs (%) | 97.3 | 97.1 | 97.2 | 97.0 | 97.1 | 97.1 | 97.0 | 96.6 | 96.2 | 96.3 | 95.9 | 96.9 | 97.2 | 97.0 | 97.0 |
| PSAURON Score (%) | 93.9 | 93.9 | 93.9 | 93.7 | 93.9 | 93.9 | 93.7 | 93.6 | 93.7 | 93.7 | 93.8 | 93.7 | 93.9 | 93.8 | 93.8 |

### Annotated gene model comparisons with B73

The 1-to-1 syntenic orthologs between B73 and each genotype identified 27,606 - 30,517 pairs, where 69.4–76.8% of B73 genes had an ortholog in any one genotype (Figure 4 & Table 3). FBLL (76.8%) and LH244 (76.1%) had the most orthologs, while LH287 (69.4%), PH2MW (69.9%), and PHP02 (70.2%) had the lowest number of orthologs. Between 10,160 and 13,277 genes per genotype had no one-to-one syntenic B73 partner, potentially because: 1) the 1-to-1 requirement leaves out tandem and other duplicate copies; 2) some genes may have moved out of syntenic position; 3) genes are derived from transposable elements; and 4) gene models were not present in the B73 annotation. As an independent check, we used Liftoff and matched each gene model to B73 genes by CDS overlap. Matching genes assigned by Liftoff agreed with MCScan 94.5–97.4% of the time (Supplementary Table 5). Liftoff matched a higher proportion of genes to B73 than MCScan (76.3–79.8%) per genotype and left fewer genes unmatched (8,730–9,950). This difference is likely because Liftoff does not require one-to-one matches; therefore, several genes can be assigned to the same B73 gene. For instance, Liftoff mapped 17–29% of genes lacking a syntenic partner to B73, suggesting that many of these genes are not absent from B73 but may potentially represent duplicated copies excluded by MCScan’s one-to-one requirement. Together, the two approaches characterize how annotated genes correspond to existing B73 annotations. MCScan identifies strict one-to-one syntenic orthologs, while Liftoff shows where a greater proportion of genes map, including duplicates. Supplementary Tables 6 and 7 provide the corresponding gene models for each B73 gene across the genotypes, as identified by MCScan (1-to-1 relationships) and Liftoff (1-to-many relationships), respectively.

**Figure 4:**
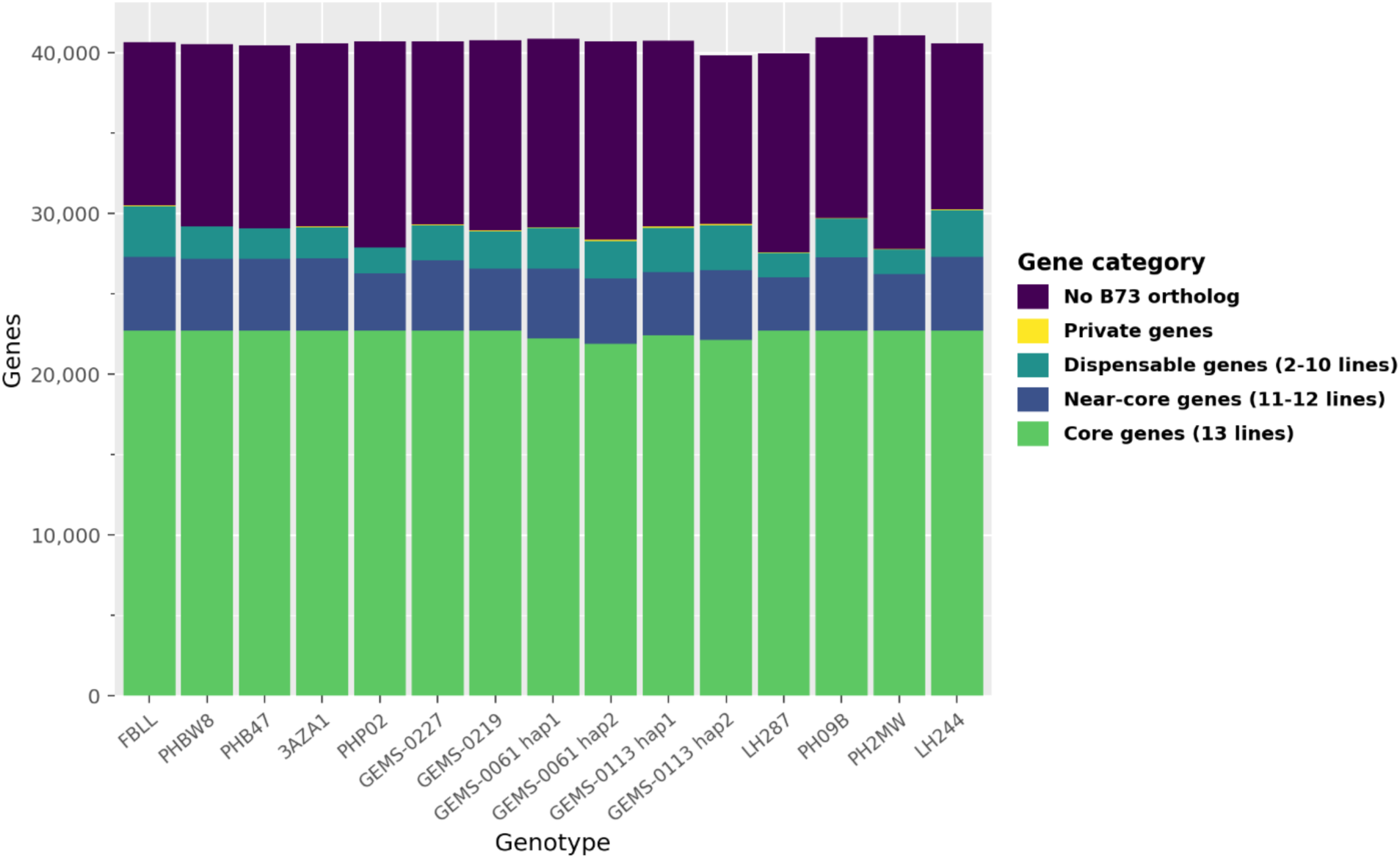
A classification of annotated genes by their similarity to existing B73 annotations (AGPv5). The core genes (green) represent genes shared across all 13 lines; near-core (blue) genes are shared in 11-12 lines; dispensable genes (teal) represent genes shared in 2-10 genotypes; private genes (yellow) represent genes found in one line; and purple represents genes that did not match any B73 gene model annotation.

**Table 3:**
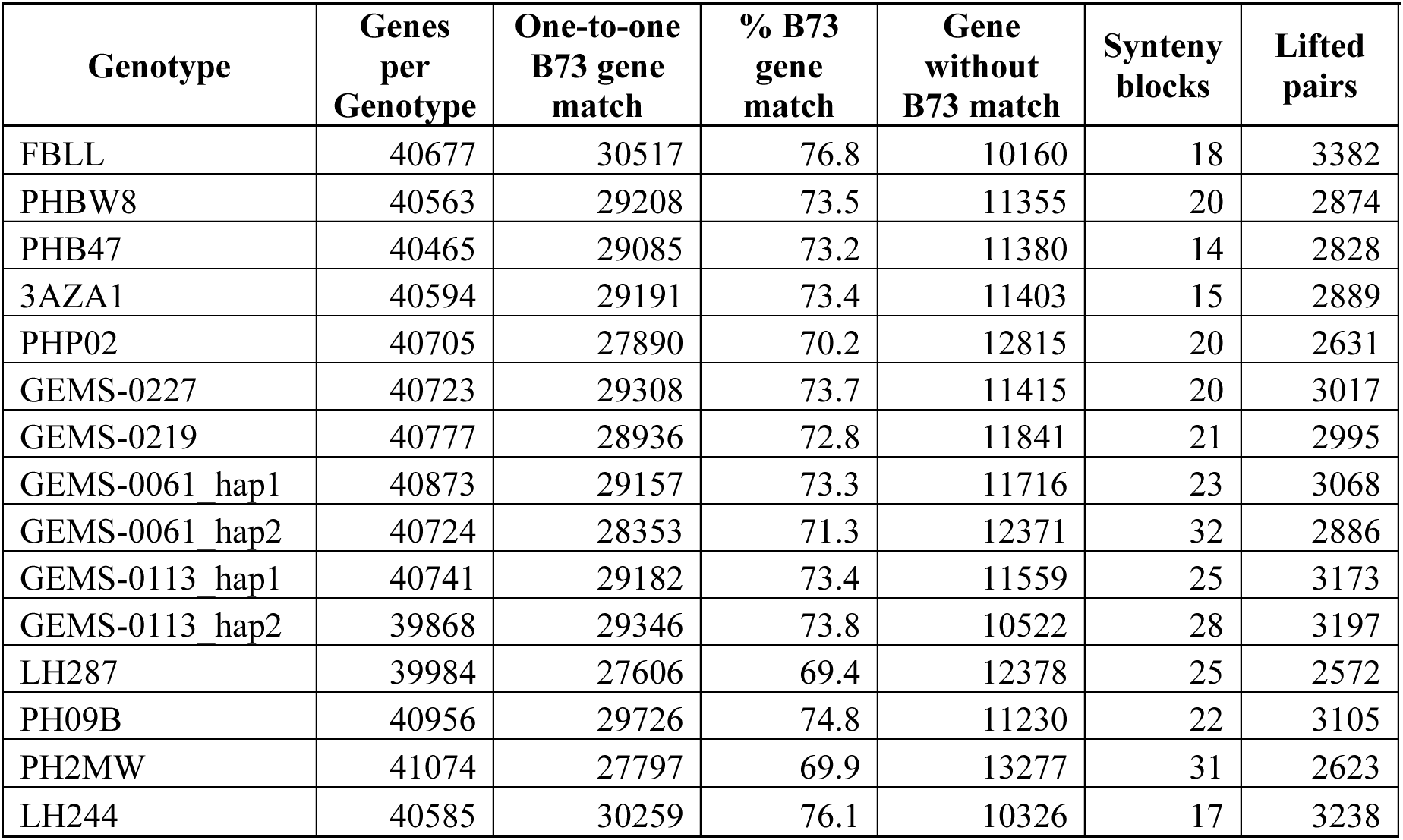
One-to-one syntenic orthologs between B73 and each genotype.

### TE Annotation

TEs occupied 1,712–2,040 Mb of the nuclear genome, corresponding to 83.6–88.6% of the genome (Supplementary Table 8), consistent with the high repeat content of maize genomes (Hufford et al., 2021). LTR retrotransposons were the most abundant class of TEs, occupying 76.9–79.6% of the nuclear genome. Structural annotation identified 54,183–59,741 intact LTR retrotransposons per genome, of which 52% were Gypsy, 38% Copia, and 11% unclassified. Intact TIR elements and intact Helitrons were collapsed into non-redundant libraries for each genome, with an average of approximately 10,000 LTR, 5,000 TIR, 2,000 Helitrons, and 16 LINE families per genome (Supplementary Table 8). Helitron detection failed in LH287, PH09B, and PH2MW; therefore, Helitron content was not reported for these genotypes, resulting in a slight underestimation of their total TE content.

### Conclusions

We present high-quality genome assemblies and annotations for diverse maize germplasm. Comparative analyses revealed shared and accession-specific structural variation, including large chromosomal inversions, highlighting genomic diversity not captured by a single reference genome. Together, these resources provide a foundation for investigating the inheritance and functional consequences of genomic variation and for improving our understanding of genotype–phenotype relationships across environments in maize.

## Data availability

Raw sequencing reads are available through the European Nucleotide Archive (ENA) under accession PRJEB82638. Genome assemblies are available through MaizeGDB under the following assembly identifiers:

- **PHBW8:** Zm-PHWB8-DRAFT-G2F-1.0
- **FBLL:** Zm-FBLL-DRAFT-G2F-1.0
- **3AZA1:** Zm-3AZA1-DRAFT-G2F-1.0
- **PHB47:** Zm-PHB47-DRAFT-G2F-1.0
- **PHP02:** Zm-PHP02-DRAFT-G2F-1.0
- **LH287:** Zm-LH287-DRAFT-G2F-1.0
- **GEMS-0113:** Zm-GEMS_0113-DRAFT-G2F-1.0
- **GEMS-0061:** Zm-GEMS_0061-DRAFT-G2F-1.0
- **GEMS-0219:** Zm-GEMS_0219-DRAFT-G2F-1.0
- **GEMS-0227:** Zm-GEMS_0227-DRAFT-G2F-1.0

## Code availability

All computational methods for assembly are documented and scripted for public use on GitHub (https://github.com/maize-genetics/g2f_2024).

## Acknowledgements

We acknowledge the Cornell BioHPC (RRID:SCR_021757) for computational resources used in this work.

## Author contributions

C.E.G., E.S.B., M.H., and M.C.R. conceived the project. N.A.J., C.E.G., and T.P.M. assembled genomes. A.M. annotated genomes. N.A.J. and A.M. conducted comparative genomics analyses. All authors contributed to manuscript drafting and revisions.

## Supplementary data

**Supplementary Table 1:** PacBio HiFi and Oxford Nanopore sequencing inputs for thirteen maize lines.

| **Genotype** | **HiFi (Gb)** | **HiFi coverage (×)** | **ONT ≥50 kb (Gb)** | **ONT coverage (×)** |
|---|---|---|---|---|
| 3AZA1 | 150.05 | 65.2 | 0.216 | 0.09 |
| FBLL | 97.03 | 42.2 | 0.974 | 0.42 |
| GEMS-0061 | 79.36 | 34.5 | 0.187 | 0.08 |
| GEMS-0113 | 74.19 | 32.3 | 0.049 | 0.02 |
| GEMS-0219 | 57.05<sup>a</sup> | 24.8 | –<sup>b</sup> | –<sup>b</sup> |
| GEMS-0227 | 103.51 | 45.0 | 0.048 | 0.02 |
| LH244 | NA | NA | NA | NA |
| LH287 | 115.74<sup>c</sup> | 50.3 | –<sup>c</sup> | –<sup>c</sup> |
| PH09B | 82.76<sup>c</sup> | 36.0 | –<sup>c</sup> | –<sup>c</sup> |
| PH2MW | 69.31<sup>c</sup> | 30.1 | –<sup>c</sup> | –<sup>c</sup> |
| PHB47 | 69.68 | 30.3 | 0.032 | 0.01 |
| PHBW8 | 84.31 | 36.7 | 0.742 | 0.32 |
| PHP02 | 75.87 | 33.0 | 1.952 | 0.85 |

*Sequencing amounts represent reads used as assembly inputs, as verified from assembly provenance or reported in collaborator documentation. ONT inputs consist of reads ≥50 kb. Coverage assumes a genome size of 2.3 Gb for both technologies. Inputs for partially phased accessions GEMS-0061 and GEMS-0113 are reported once per accession. NA indicates unavailable sequencing records; – indicates no ONT assembly input documented.*
<sup>a</sup> *HiFi reads from a single SMRT cell; two additional cells, derived from a different plant of the same accession, were excluded.*
<sup>b</sup> *ONT reads were generated but excluded from assembly because their plant of origin relative to the HiFi cell used could not be verified.*
<sup>c</sup> *HiFi quantities are from collaborator-provided read sets. Provider documentation describes HiFi-only assemblies, although ONT reads were also generated. ONT usage remains to be independently confirmed.*

**Supplementary Table 2:** Summary statistics for the genomic features identified.

| **Genotype** | **Mean CDS length (bp)** | **Mean Protein length (aa)** | **Mean Gene length (bp)** | **Mean Exons per gene** |
|---|---|---|---|---|
| FBLL | 1136.5 | 377.8 | 4276.9 | 5.88 |
| PHBW8 | 1145.2 | 380.7 | 4290.7 | 5.9 |
| PHB47 | 1138.5 | 378.5 | 4291.9 | 5.88 |
| 3AZA1 | 1136.2 | 377.7 | 4278.8 | 5.89 |
| PHP02 | 1138.8 | 378.6 | 4294.6 | 5.9 |
| GEMS-0227 | 1137.2 | 378.1 | 4271.4 | 5.89 |
| GEMS-0219 | 1138.1 | 378.4 | 4272.5 | 5.89 |
| GEMS-0061_hap1 | 1134.8 | 377.3 | 4253.1 | 5.86 |
| GEMS-0061_hap2 | 1134.4 | 377.1 | 4265.8 | 5.87 |
| GEMS-0113_hap1 | 1140.9 | 379.3 | 4286 | 5.9 |
| GEMS-0113_hap2 | 1141.6 | 379.5 | 4302.8 | 5.91 |
| LH287 | 1140.6 | 379.2 | 4293.8 | 5.88 |
| PH09B | 1138.5 | 378.5 | 4263.7 | 5.87 |
| PH2MW | 1139 | 378.7 | 4273.7 | 5.87 |
| LH244 | 1137.7 | 378.2 | 4268.7 | 5.88 |

**Supplementary Table 3.** BUSCO completeness of the Helixer gene annotations.

| **Genotype** | **Complete (C, %)** | **Single-copy (S, %)** | **Duplicated (D, %)** | **Fragmented (F, %)** | **Missing (M, %)** |
|---|---|---|---|---|---|
| FBLL | 97.3 | 72.2 | 25.1 | 1.3 | 1.4 |
| PHBW8 | 97.1 | 71.7 | 25.4 | 1.3 | 1.5 |
| PHB47 | 97.2 | 72.3 | 24.9 | 1.3 | 1.5 |
| 3AZA1 | 97 | 72 | 25 | 1.4 | 1.6 |
| PHP02 | 97.1 | 71.7 | 25.4 | 1.4 | 1.4 |
| GEMS-0227 | 97.1 | 70.1 | 27 | 1.3 | 1.5 |
| GEMS-0219 | 97 | 71.1 | 25.9 | 1.2 | 1.8 |
| GEMS-0061 hap1 | 96.6 | 70.4 | 26.2 | 1.3 | 2.1 |
| GEMS-0061 hap2 | 96.2 | 70.1 | 26.1 | 1.4 | 2.4 |
| GEMS-0113 hap1 | 96.3 | 70.5 | 25.8 | 1.3 | 2.4 |
| GEMS-0113 hap2 | 95.9 | 71.1 | 24.8 | 1.2 | 2.8 |
| LH287 | 96.9 | 71.9 | 25 | 1.5 | 1.6 |
| PH09B | 97.2 | 71.7 | 25.5 | 1.2 | 1.6 |
| PH2MW | 97 | 71.8 | 25.3 | 1.3 | 1.6 |
| LH244 | 97 | 72.1 | 24.9 | 1.4 | 1.6 |

**Supplementary Table 4:**
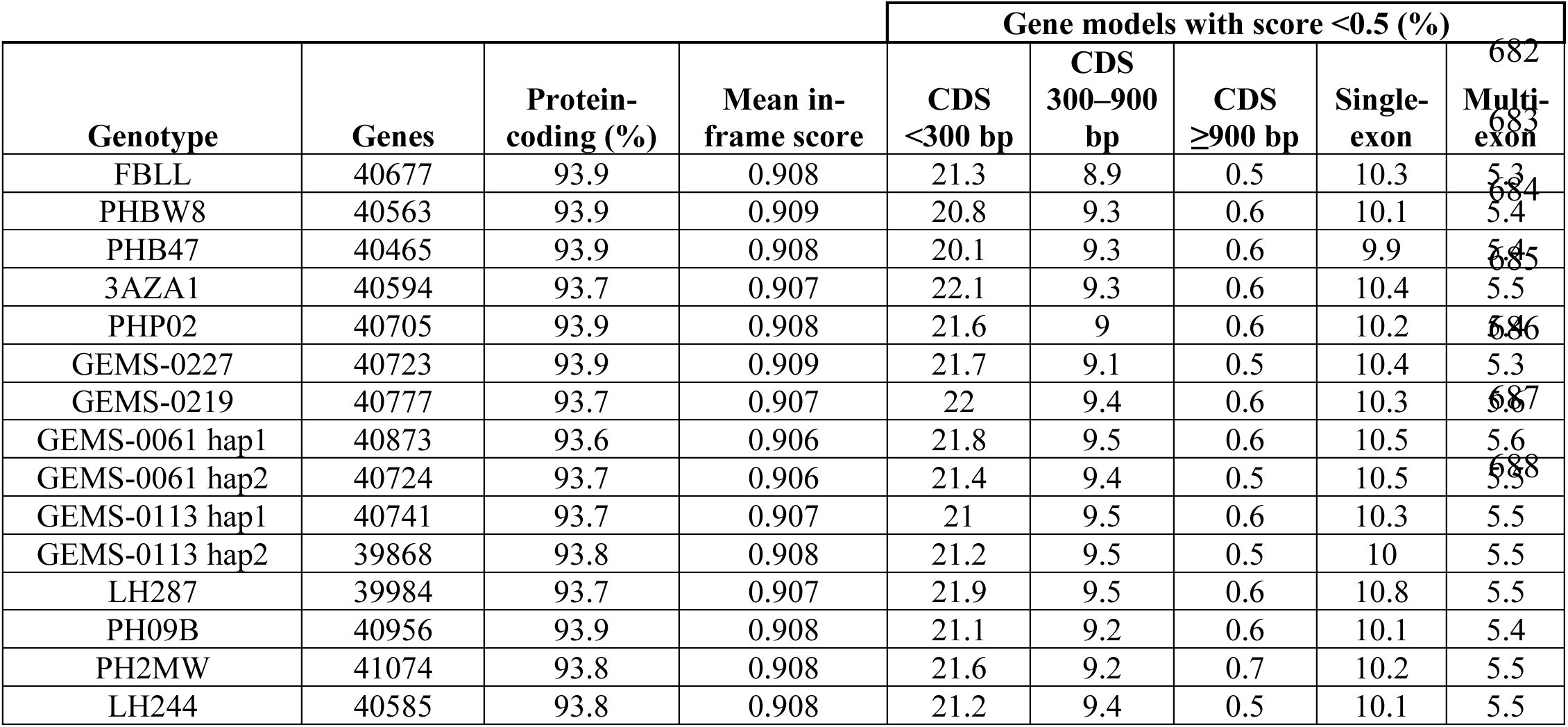
Summary statistics of PSAURON assessment of predicted gene models across the genotypes.

**Supplementary Table 5:** Summary of gene model comparisons to B73 across the genotypes using Liftoff.

| **Genotype** | **Genes per Genotype** | **Genes matches** | **% gene matched** | **Genotype genes matched** | **Genes without B73 match** | **B73 genes with one match** | **B73 genes with ≥2 matches** | **Genes matched to ≥2 B73 genes** |
|---|---|---|---|---|---|---|---|---|
| FBLL | 40677 | 31155 | 79.8 | 31947 | 8730 | 30069 | 1086 | 464 |
| PHBW8 | 40563 | 30439 | 78 | 31205 | 9358 | 29407 | 1032 | 427 |
| PHB47 | 40465 | 30407 | 77.9 | 31188 | 9277 | 29357 | 1050 | 422 |
| 3AZA1 | 40594 | 30534 | 78.2 | 31261 | 9333 | 29532 | 1002 | 431 |
| PHP02 | 40705 | 30337 | 77.7 | 31061 | 9644 | 29312 | 1025 | 430 |
| GEMS-0227 | 40723 | 29990 | 76.8 | 30773 | 9950 | 28946 | 1044 | 421 |
| GEMS-0219 | 40777 | 30637 | 78.5 | 31385 | 9392 | 29560 | 1077 | 471 |
| GEMS-0061_hap1 | 40873 | 30426 | 77.9 | 31213 | 9660 | 29368 | 1058 | 431 |
| GEMS-0061_hap2 | 40724 | 30265 | 77.5 | 31065 | 9659 | 29217 | 1048 | 416 |
| GEMS-0113_hap1 | 40741 | 30482 | 78.1 | 31225 | 9516 | 29428 | 1054 | 463 |
| GEMS-0113_hap2 | 39868 | 30251 | 77.5 | 30974 | 8894 | 29228 | 1023 | 448 |
| LH287 | 39984 | 29792 | 76.3 | 30493 | 9491 | 28811 | 981 | 403 |
| PH09B | 40956 | 30729 | 78.7 | 31441 | 9515 | 29704 | 1025 | 460 |
| PH2MW | 41074 | 30408 | 77.9 | 31128 | 9946 | 29391 | 1017 | 425 |
| LH244 | 40585 | 30977 | 79.4 | 31764 | 8821 | 29883 | 1094 | 479 |

**Supplementary Tables 6 & 7.** in attached document.

**Supplementary Table 8:** Summary statistics for all the TE features identified.

| **Genotype** | **All TEs (% )** | **LTR: intact elements** | **LTR: families** | **LTR: content (%)** | **TIR: intact elements** | **TIR: families** | **Helitron: intact elements** | **Helitron: families** | **LINE: families** | **SINE: families** |
|---|---|---|---|---|---|---|---|---|---|---|
| FBLL | 88.28 | 58,727 | 10,360 | 78.98 | 21,892 | 4,972 | 4,357 | 2,312 | 17 | 0 |
| PHBW8 | 88.23 | 58,576 | 10,684 | 79.09 | 21,732 | 5,227 | 4,393 | 2,303 | 15 | 0 |
| PHB47 | 87.02 | 58,254 | 10,513 | 78.46 | 21,435 | 5,120 | 4,417 | 1,528 | 16 | 0 |
| 3AZA1 | 88.61 | 59,741 | 10,119 | 79.59 | 21,832 | 5,359 | 4,398 | 2,356 | 17 | 0 |
| PHP02 | 87.89 | 58,665 | 9,840 | 78.79 | 20,516 | 4,844 | 4,455 | 1,932 | 19 | 0 |
| GEMS-0227 | 87.53 | 59,635 | 10,219 | 78.53 | 21,767 | 5,149 | 4,366 | 1,991 | 14 | 0 |
| GEMS-0219 | 87.78 | 59,143 | 10,667 | 78.29 | 21,549 | 5,049 | 4,404 | 2,238 | 10 | 0 |
| GEMS-0061 hap1 | 86.93 | 59,392 | 8,752 | 78.13 | 21,093 | 5,218 | 4,470 | 1,931 | 19 | 0 |
| GEMS-0061 hap2 | 88.2 | 58,394 | 10,729 | 79.31 | 21,269 | 5,180 | 4,451 | 2,175 | 18 | 0 |
| GEMS-0113 hap1 | 88.08 | 59,045 | 10,501 | 78.76 | 21,786 | 4,863 | 4,408 | 2,148 | 21 | 0 |
| GEMS-0113 hap2 | 87.72 | 58,730 | 10,253 | 78.74 | 21,653 | 4,934 | 4,341 | 2,370 | 17 | 0 |
| LH287 | 83.63* | 54,183 | 9,529 | 76.89 | 20,982 | 4,612 | NA | NA | 12 | 0 |
| PH09B | 86.30* | 59,545 | 10,236 | 79.17 | 21,735 | 4,704 | NA | NA | 17 | 0 |
| PH2MW | 86.02* | 58,561 | 10,418 | 79.01 | 20,847 | 4,990 | NA | NA | 15 | 0 |
| LH244 | 87.44 | 59,439 | 10,695 | 78.79 | 21,879 | 5,023 | 4,418 | 1,291 | 13 | 0 |

